# Croatian Genetic Heritage: Renewed Y Chromosome Story Two Decades Later

**DOI:** 10.1101/2022.03.21.485134

**Authors:** Dragan Primorac, Vedrana Škaro, Petar Projić, Saša Missoni, Ivana Horjan Zanki, Sinisa Merkaš, Jelena Šarac, Natalija Novokmet, Andrea Ledić, Adela Makar, Gordan Lauc, Šimun Anđelinović, Željana Bašić, Ivana Kružić, Marijana Neuberg, Martina Smolić, Robert Smolić, Irena Hrstić, Dragan Trivanović, Rijad Konjhodžić, Lana Salihefendić, Naida Babić Jordamović, Damir Marjanović

**Affiliations:** St. Catherine Hospital, Zagreb, Croatia; School of Medicine, University of Split, Split, Croatia; University Department of Forensic Sciences, University of Split, Split, Croatia; Faculty of Medicine, University of Osijek, Osijek, Croatia; Faculty of Dental Medicine and Health, University of Osijek, Osijek Croatia; School of Medicine Rijeka, University of Rijeka, Croatia; Eberly College of Science, Pennsylvania State University, University Park, PA, USA; Henry C. Lee College of Criminal Justice and Forensic Sciences, University of New Haven, West Haven, CT, USA; Medical School REGIOMED, Coburg, Germany; The National Forensic Sciences University, Gandhinagar, Gujarat, India; Molecular Anthropology Laboratory, Center for Applied Bioanthropology, Institute for Anthropological Research, Zagreb, Croatia; DNA Laboratory, Genos Ltd., Zagreb, Croatia; Department of Pharmacology, Faculty of Pharmacy and Biochemistry, University of Zagreb; Forensic Science Centre „Ivan Vučetić“, Zagreb, Croatia; University Hospital Center, Split, Croatia; University North, Varaždin, Croatia; General Hospital Pula, Pula, Croatia; Alea Genetic Center, Sarajevo, Bosnia and Herzegovina; Department of Genetics and Bioengineering, International Burch University, Sarajevo, Bosnia and Herzegovina

**Author notes:** **Correspondence to**: Vedrana Škaro, Genos Ltd., DNA Laboratory, Zagreb, Croatia.

## Abstract

**Aim:** To analyze an additional set of Y-Chromosome genetic markers to acquire a more detailed insight into the diversity of the Croatian population.

**Methods:** The total number of 518 Yfiler™ Plus profiles were genotyped. Allele, haplotype frequencies, and haplotype diversity were calculated using the STRAF software package v2.0.4. Genetic distances were quantified by *R*st using AMOVA online tool from the YHRD. The evolutionary history was inferred using the neighbor-joining method of phylogenetic tree construction in MEGAX software. Whit Athey’s Haplogroup Predictor v5 was used for additional comparison with available regional and other European populations.

**Results:** The total of 507 haplotypes were used for genetic STR analysis. The interpopulation study on 17 Y-STR markers shows the lowest genetic diversity between the Croatian and Bosnian-Herzegovinian populations and the highest between the Croatian and Irish populations. Additional interpopulation comparison with the original 27 Y-STR markers (for the population with available data) was also performed. A total of 518 haplotypes were used in the determination of haplogroup diversity. Haplogroup I with its sublineage I2a expressed the highest prevalence. Haplogroup R, with its major sublineage R1a, is the second most abundant in the studied Croatian population, except for the subpopulation of Hvar, where E1b1b is the second most abundant haplogroup. Rare haplogroups also confirmed in this study are L, T, and Q. G1 is detected for the very first time in the Croatian population.

**Conclusion:** New insight into differences between examined subpopulations of Croatia and their possible (dis)similarities with neighboring abroad populations was notified.

---

The Y chromosome (~ 60 Mb) is relatively small and inherited from father to son unchanged (except for occasional mutations). Except for the small pseudoautosomal regions (PAR), there is no recombination between the X and Y chromosome (1–3). This event is the reason why haplotype inheritance through the male lineage can be tracked and analyzed (2, 4–6).

The Y chromosome mostly consists of repetitive sequences (around 50%), which are single-base substitutions, Alu elements, and Long Interspersed Nuclear Elements (LINEs). Short tandem repeats (STRs), as repetitive elements are the base of population genetic studies. Their average mutational frequency is ~0.2% per generation (7, 8).

Y haplogroup can be defined as a part of the Y chromosome family related by ancestry and determined by a specific set of Y chromosomal single nucleotide polymorphisms (Y-SNPs). It is of great importance to better understand of the demographic processes that shaped modern populations (8, 9). The low mutation rate makes Y-SNP markers suitable for the conventional method of Y haplogroup defining.

Y chromosome haplogroups can also be successfully predicted from Y-STR markers (Y-STR haplotype) using Y-STR haplogroup predicting tools. Lately, this method has gained much attention due to its labor-, time-, and cost-effectiveness (10). The haplotype helps analyze the influence of genes on disease-related alleles and represents the set of alleles on the same chromosome. On the other hand, major haplogroups (branches of Y-chromosome phylogeny), labeled A-T, define the establishment and expansion of major population groups and can indicate the time scale and the route of major migration events (11).

The main aim of this research is to update information about Croatian Y chromosome diversity by using additional Y STR loci to compare new results with the previously published results generated using Y-STR and Y-SNP markers (12–20).

Another goal was to analyze the genetic structure of five regional subpopulations (with the local centers in Osijek, Pula, Varaždin, Split, and Hvar Island) by identifying the most common haplogroups in these regions. The analysis also included genetic differences between these five subpopulations and their potential (dis)similarity with neighboring countries.

## Materials and methods

For this study, buccal swab samples were obtained from 518 unrelated adult male individuals from five different regions of Croatia: Hvar (n=104), Osijek (n=110), Pula (n=99), Varaždin (n=100), and Split (n=105).

DNA extraction was performed on QIAsymphony instrument using the QIAsymphony DNA Investigator Kit and protocol (Qiagen, Hilden, Germany). DNA was quantified on Rotor-Gene Q real-time PCR cycler (Qiagen, Hilden, Germany) using Q-Rex Software and Investigator Quantiplex Pro RGQ Kit. Yfiler™ Plus PCR Amplification Kit (Applied Biosystems, Foster City, CA, USA) was used to simultaneously amplify 27 Y chromosome STR loci. Amplification was carried out following the manufacturer’s protocol. PCR amplification was performed on Mastercycler® nexus SX1 PCR thermal cycler (Eppendorf AG, Hamburg, Germany) according to the manufacturer’s instructions. PCR-amplified products were separated and detected using standard protocols for electrophoresis on 3500 Genetic Analyzer (Applied Biosystems, Foster City, CA, USA). Allele calling was performed with GeneMapper™ ID-X Software v1.4 (Applied Biosystems, Foster City, CA, USA) using the custom panel and bin sets.

A total number of 507 fully genotyped Y-STR profiles of the present study were submitted to Y Chromosome Haplotype Reference Database (YHRD) with the accession numbers assigned as follows: Hvar (n=104; YA004742), Osijek (n=109; YA004743), Pula (n=94; YA004744), Varaždin (n=98; YA004746), and Split (n=102; YA004745).

### Statistical analysis

The number of alleles and different haplotypes, allele and haplotype frequencies, and gene and haplotype diversity were estimated in order to assess the intrapopulation diversity.

Haplotype diversity was calculated using Nei’s formula: HD = (1-∑*p_i_*^2^)*n/(n-1), where n is the sample size and *p_i_* is the *i*^th^ haplotype frequency. Gene diversity was calculated as 1-∑*p_i_*^2^, where *p_i_* is the allele frequency. The formula ∑*p_i_*^2^ was used to calculate match probability (MP), where *p_i_* is the frequency of the *i*^th^ haplotype. Discrimination capacity (DC) was calculated by dividing the number of haplotypes by the number of individuals in the population (21, 22). Allele and haplotype frequencies, as well as gene and haplotype diversity, were calculated using the STRAF software package v2.0.4 (23, 24).

Genetic distances between groups of males and between populations were quantified by *R*_*s*t_ using AMOVA online tool from the YHRD (25, 26). In addition, associated probability values (*P* values) with 10,000 permutations were included for the studied populations. Genetic distances were used to generate the multidimensional scaling plot (MDS) plots for the comparison of population haplotype data from YHRD.

AMOVA analysis was performed with two population groups. The number of the populations with available data for 27 STR loci was relatively small, especially in the closest Croatian neighborhood. Therefore, the first group has been analyzed by comparing the reduced number of markers using the 17 Y-STR loci included in the AmpFLSTR™ Yfiler™ PCR Amplification Kit. The second group has been analyzed by comparing the whole set of 27 Y-STR loci included in the Yfiler™ Plus PCR Amplification Kit.

The first group of European populations selected for comparison with the population of Croatia using 17 Y-STRs included: Tiroler Unterland, Austria (n=547), Antwerpen, Belgium (n=309), Bosnia and Herzegovina (n=574), Bulgaria (n=91), Rostock, Germany (n=598), Greece (n=191), Hungary (n=303), Italy (n=147), Warsaw, Poland (n=491), Serbia (n=567), Albania (n=315), Czech Republic (n=109), Estonia (n=123), Ireland (n=863), Lithuania (n=531), North Macedonia (n=493), Norway (n=1555), Slovenia (n=294), Sweden (n=296), and Ukraine (n=212).

The second group of the Worldwide populations selected for comparison with the population of Croatia using the 27 Y-STRs: Croatia (n = 507, present study), Slovenia (n = 194, Belgium (n = 160, Hungary (n = 218), Austria (n =392), Germany (n = 495), Italy (n = 689), North Macedonia (n = 295), Serbia (n = 183), Denmark (n=177), Ethiopia (n=290), French Polynesia (n=81), Ghana (n=584), India (n=541), Lithuania (n=251), Mexico (n=354), Nigeria (n=337), Pakistan (n=280), Poland (n=612), Russian Federation (n=958), Saudi Arabia (n=156), Spain (n=316,), Switzerland (n=724), and United Kingdom (n=115).

Available population data and all related references are included in the YHRD (25, 26). The evolutionary history was inferred for both sets of markers using the neighbor-joining (NJ) method of phylogenetic tree construction (27) in MEGAX (28), whereby the optimal tree is shown.

Y-chromosomal haplogroup prediction using allele frequencies on 518 Yfiler™ Plus profiles was performed using Whit Athey’s Haplogroup Predictor v5, an algorithm based on the Bayesian-allele-frequency approach (29, 30).

## Results and discussion

A total of 518 haplotypes were detected and used for haplogroup prediction. Eleven haplotypes were considered newly detected microvariants, which require additional analysis for confirmation. Therefore, the remaining 507 haplotypes (the ones without newly detected microvariants) were used for additional statistical analysis. On a sample of fully genotyped 507 Y-STR profiles, a total of 502 different haplotypes were detected in the study, with 497 unique haplotypes and 5 haplotypes appearing twice. In addition, 196 alleles at 27 Y-STR loci were detected (Table 1). Apart from the DYS385a/b double locus, the largest number of alleles was recorded on DYS481 with 14 detected alleles. Three loci had the smallest number of alleles, namely DYS393, DYS437, and Y-GATA-H4 with four alleles each.

**TABLE 1.**
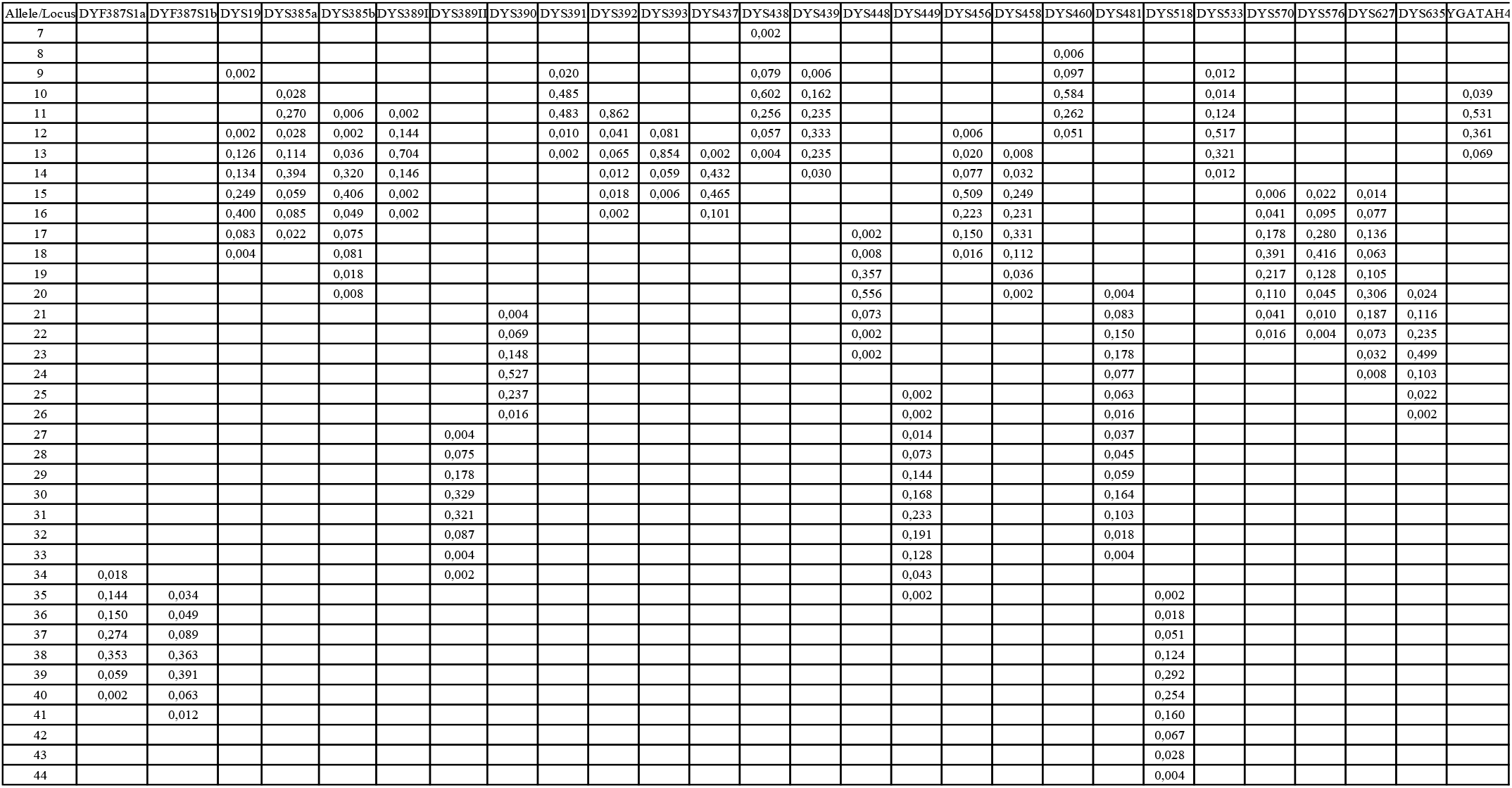
Allele frequencies for the 27 Y-STR loci in the population of Croatia (n=518). The study included five Croatian regional subpopulations with the local centers in Osijek, Pula, Varaždin, Split, and Hvar Island. Yfiler™ Plus PCR Amplification Kit (Applied Biosystems) was used.

Haplotype diversity was calculated to be 1.0000 ± 0.0014 with DC of 1.00 and MP of 0.01. The average genetic diversity for the study population was 0.656 across all loci, ranging from 0,886 for DYS481to 0,251 for DYS392. At the population level, the most common allele is allele 11 at locus DYS392 with a frequency of 0.862. This was not surprising considering that DYS392 is one of the least polymorphic loci in the studied population with six detected alleles and the lowest genetic diversity.

In order to determine additional genetic differences within the analyzed population of Croatia, an interpopulation analysis was done between five regions: Hvar (n=104), Osijek (n=109), Pula (n=94), Split (n=102) and Varaždin (n=98). The lowest genetic diversity observed for the population of Hvar was compared to the population of Split (*R*st=0.0009, *P*=0.3240). The greatest genetic diversity observed for the population of Hvar was compared to the population of Varaždin (*R*st=0.0979, *P*=0.0000), the population of Pula (*R*st=0.0284, *P*=0.0042), and the population of Osijek (*R*st=0.0210, *P*=0.0097). The lowest genetic diversity observed for the population of Osijek was compared to the population of Split (*R*st=0.0063, *P*=0.1199) and the population of Pula (*R*st=0.0069, *P*=0.1138). The greatest genetic diversity observed for the population of Osijek was compared to the population of Varaždin (*R*st=0.0551, *P*=0.0000) and the population of Hvar (*R*st=0.0210, *P*=0.0097). The lowest genetic diversity observed for the population of Pula was compared to the population of Osijek (*R*st=0.0069, *P*=0.1138). The greatest genetic diversity observed for the population of Pula was compared to the population of Hvar (*R*st==0.0284, *P*=0.0042), the population of Split (*R*st=0.0180, *P*=0.0233) and the population of Varaždin (*R*st=0.0166, *P*=0.0260). The lowest genetic diversity observed for the population of Split was compared to population of Hvar (*R*st=0.0009, *P*=0.3240) and the population of Osijek (*R*st=0.0063, *P*=0.1199). The greatest genetic diversity observed for the population of Split was compared to the population of Varaždin (*R*st=0.0821, *P*=0.0000) and the population of Pula (*R*st=0.0180, *P*=0.0233). The lowest genetic diversity observed for the population of Varaždin was compared to the population of Pula (*R*st=0.0166, *P*=0.0260). The greatest genetic diversity observed for the population of Varaždin was compared to the population of Hvar (*R*st=0.0979, *P*=0.0000), the population of Split (*R*st=0.0821, *P*=0.0000), and the population of Osijek (*R*st=0.0551, *P*=0.0000).

In order to compare the studied population with a large number of worldwide published population data, interpopulation analyses were completed by comparing the analyzed population with two groups of countries.

The first group of populations selected for comparison with the population of Croatia using reduced 17 Y-STR set of markers included 20 populations. The lowest genetic diversity was observed between the currently analyzed population of Croatia and previously published population of Bosnia and Herzegovina (*R*_st_=0.0076, *P*=0.0002), and the population of Serbia (*R*_st_= 0.0186, *P*=0.000). Other populations with low genetic diversity values when compared to the present results include those from Bulgaria (*R*_st_=0.0144, *P*=0.000), Ukraine (*R*st=0.0195, *P*=0.000), Slovenia (*R*_st_=0.0204, *P*=0.000), Hungary (*R*_st_=0.0238, *P*=0.0000), Greece (*R*_st_=0.0241, *P*=0.0000), North Macedonia (*R*_st_=0.0375, *P*=0.0000), Italy (*R*_st_=0.0659, *P*=0.0000), Albania (*R*_st_=0.0728, *P*=0.0000), Czech Republic (*R*_st_=0.0767, *P*=0.000), and Austria (*R*_st_=0.0795, *P*=0.0000). The highest genetic distance was observed when the study population was compared with the populations of Ireland (*R*_st_= 0.3178, *P*=0.000), Estonia (*R*_st_=0.1877, *P*=0.0000), Lithuania (*R*_st_=0.1706, *P*=0.0000), Belgium (*R*_st_=0.1429, *P*=0.0000), Norway (*R*_st_=0.1270, *P*=0.0000) Poland (*R*_st_=0.1216, *P*=0.0000), Sweden (*R*_st_=0.1209, *P*=0.0000), and Germany (*R*_st_=0.1036, *P*=0.0000).

The second group of selected countries used the set of all 27 Y-STR markers. The selection was limited since this is an expanded panel of Y-STR markers, and data are not available for many populations.

By comparing the population from the present study with previously published data of the first selected group on 27 Y-STR markers for 23 populations, the lowest genetic diversity was observed between the currently analyzed population of Croatia and the previously published Serbian population (*R*st=0.0097, *P*=0.0055), and the population of Slovenia (*R*st= 0.0297, *P*=0.000). Other populations with low genetic diversity values when compared to the present results include those from Hungary (*R*st=0.0482, *P*=0.000), North Macedonia (*R*st=0.0720, *P*=0.0000), Russian Federation (*R*st=0.0779, *P*=0.0000), Pakistan (*R*st=0.0854, *P*=0.0000), Poland (*R*st=0.0905, *P*=0.0000), India (*R*st=0.0961, *P*=0.0000), and Saudi Arabia (*R*st=0.0990, *P*=0.0000). The highest genetic distance was observed when the study population was compared with the populations of Spain (*R*st= 0.3283, *P*=0.000), Ghana (*R*st=0.2787, *P*=0.0000), Nigeria (*R*st=0.2414, *P*=0.0000), Italy (*R*st=0.2043, *P*=0.0000), Belgium (*R*st=0.2031, *P*=0.0000), Switzerland (*R*st=0.1999, *P*=0.0000), Lithuania (*R*st=0.1914, *P*=0.0000), French Polynesia (*R*st=0.1878, *P*=0.0000), Mexico (*R*st=0.1785, *P*=0.0000), Ethiopia (*R*st=0.1704, *P*=0.0000), Germany (*R*st=0.1461, *P*=0.0000), Denmark (*R*st=0.1290, *P*=0.0000), Austria (*R*st=0.1255, *P*=0.0000), and United Kingdom (*R*st=0.1122, *P*=0.0000).

To further investigate molecular evolutionary relationships between the geographical subpopulations of Croatia, NJ phylogenetic trees were constructed based on *R*st values for different regions of Croatia (Figure 1).

**FIGURE 1.**
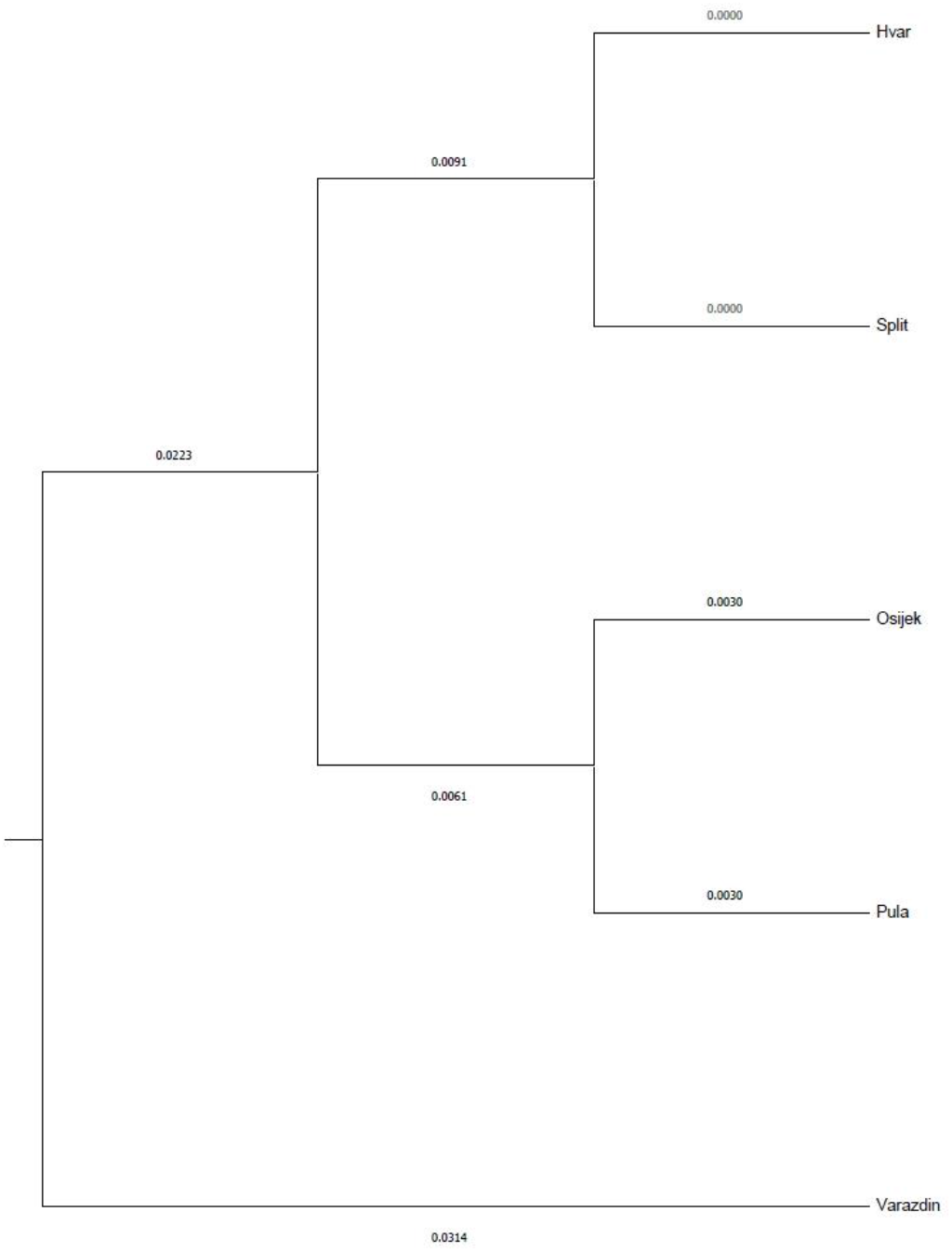
The neighbor-joining phylogenetic tree shows the genetic relationships and clustering between five Croatian regions based on the population study of 27 Y-STR markers (Yfiler™ Plus PCR Amplification Kit, Applied Biosystems).

Genetic relationships between investigated populations are shown in MDS plots (Figures 2 and 3). The results of such comparisons confirm the general trends that were shown in Supplementary Tables 1 and 2.

**FIGURE 2.**
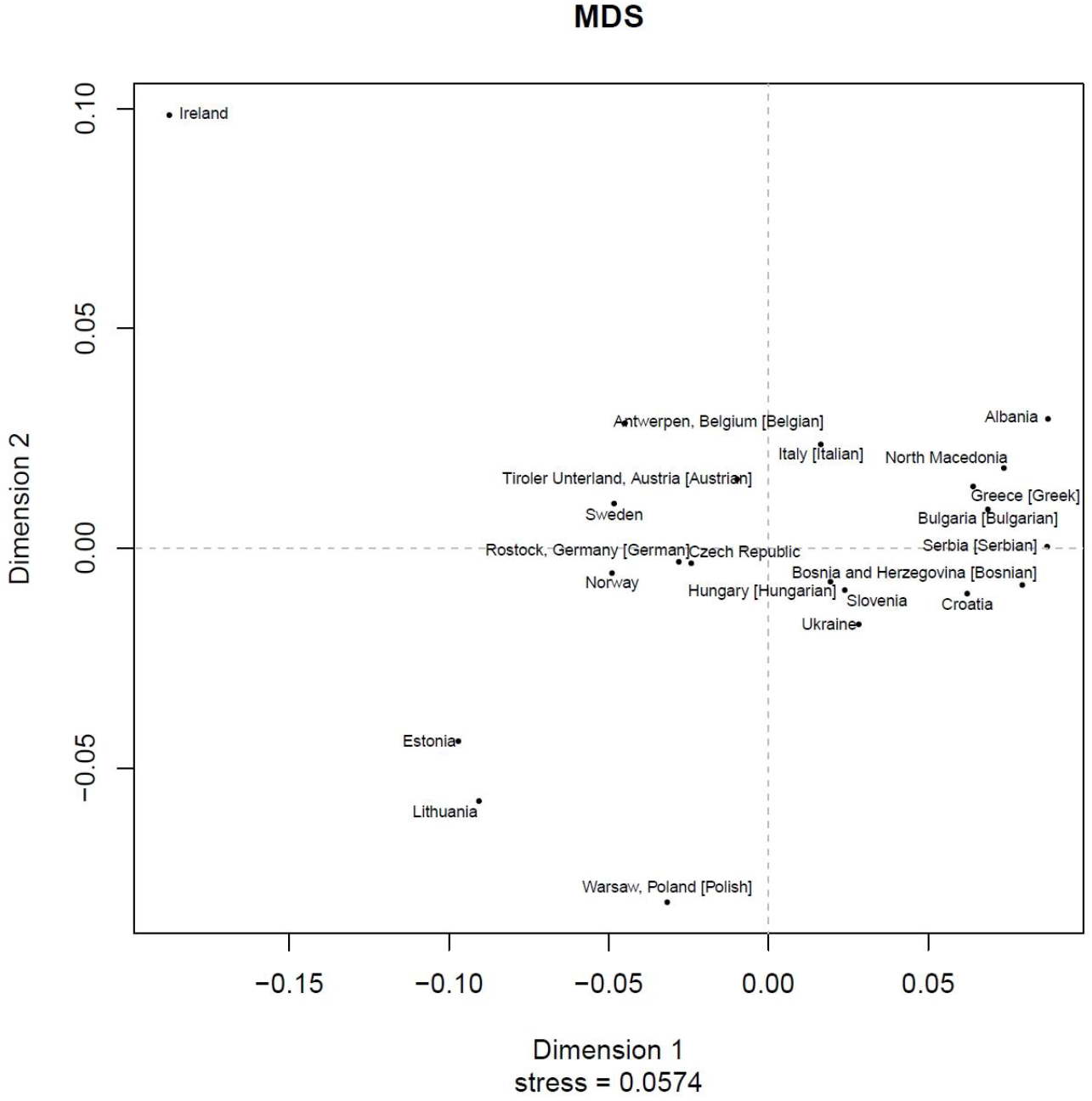
MDS plot showing genetic differentiation between the 21 populations in two dimensions, based on analysis of available data for 17 Y-STR markers included in the Yfiler™ marker set (Applied Biosystems).

**FIGURE 3.**
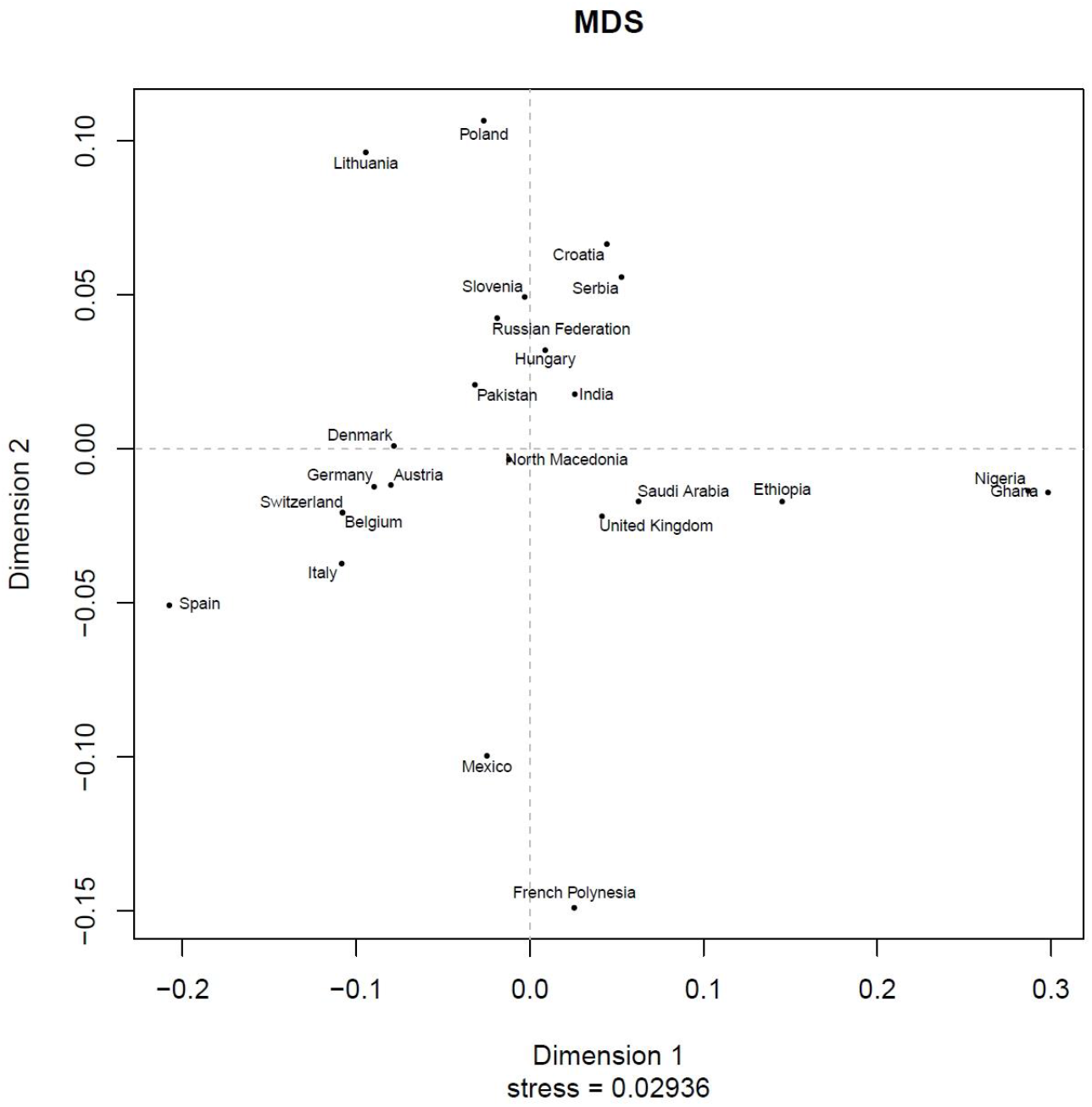
MDS plot showing genetic differentiation between 24 analyzed populations in two dimensions, based on analysis of available data for 27 Y-STR markers included in the Yfiler™ Plus PCR Amplification Kit (Applied Biosystems). Data for some neighboring populations (i.e., Bosnia and Herzegovina and Monte Negro) and a few others that showed clustering when analyzed on 17 Y-STR markers (e.g., Bulgaria, Albania, Ukraine etc.) were not available and therefore not shown.

The NJ phylogenetic tree shows the genetic relationships and clustering between five Croatian regions based on the population study on 27 Y-STR markers (Figure 1). Hvar and Split subpopulations are clustered together. Osijek and Pula subpopulations are in a separate cluster. The population of the Varaždin region is in a cluster on a different branch that may indicate its genetic specificity (probably linked with geographical position) relative to the other four examined regional subpopulations.

In the MDS plot showing the group of populations compared using the set of 17 Y-STR markers, as expected, the Croatian population is clustered with the geographically close populations of Bosnia and Herzegovina, Serbia, and Slovenia. The populations of Ukraine, Bulgaria, and Hungary are also clustered close to those four populations. The next closest European populations are North Macedonia, Albania, and the Czech Republic (Figure 2).

In the MDS plot showing the group of populations compared using the set of 27 Y-STR markers, the Croatian population is closely clustered with the geographically close populations of Serbia and Slovenia. The populations of Poland, Hungary, and the Russian Federation are also clustered relatively close to those three populations. The next closest European population is North Macedonia.

(Figure 3). A comparison between the Croatian population and some neighboring populations (i.e., Bosnia and Herzegovina and Monte Negro) and a few others that showed clustering when analyzed on 17 Y-STR markers (e.g., Bulgaria, Albania, Ukraine etc. was not performed since data on 27 Y-STR markers are unavailable for those populations.

Our study showed a high degree of homogeneity of the Croatian population. Certain genetic similarity has been observed at the regional level (between the the population of the Pula region and Serbian population (*R*st= 0.0063, *P*=0.1013), and between the population of the Varaždin region and neighboring Slovenian population (*R*st= = −0.0002, *P*= 0.4124)). These results could prove again that the Y-chromosome is expected to show greater geographical clustering than other population markers (2, 16), but also could potentially mark immigrational impacts from the eastern neighboring countries, such as those in the Istrian region, most probably in the second half of the 20^th^ century. However, these initially notified similarities still should be confirmed by additional analysis and increasing/structuring sample size of the Pula and Varaždin region.

For the calculation of Y-chromosomal haplogroup prediction and intrapopulation variability between the five subpopulations, the total number of 518 Yfiler™ Plus profiles was used: Hvar (n=104), Varaždin (n=100), Split (n=105), Pula (n=99), and Osijek (n=110). Regarding the haplogroup diversity between these five subpopulations of Croatia, successful haplogroup assignment was obtained for all 518 Y-STR profiles (Table 2). The results of Y haplogroup prediction using Whit Athey’s Haplogroup Predictor tool (182) are summarized in Figure 4. Prediction accuracy was estimated to be 100% in 492 cases. For the remaining 26 samples the prediction accuracy was 5%. Prediction accuracy varied between 63,1% and 99.58%. Out of a total of 14 detected haplogroups, the most prevalent one is I2a, which accounts for 39% of all samples, followed by R1a (24,32%) and E1b1b (10,18%). The remaining eight haplogroups were less frequent.

**TABLE 2.** Haplogroup composition in the five regions of Croatia with the local centers in Osijek (n=110), Pula (n=99)., Varaždin (n=100), Split (n=105), and Hvar Island (n=104). Y chromosome haplogroups prediction is based on the population study of 27 Y-STR markers (Yfiler™ Plus PCR Amplification Kit, Applied Biosystems).

|  | Haplogroup/number of haplotypes |  |  |  |  |  |  |  |  |  |  |  |  |  |
| --- | --- | --- | --- | --- | --- | --- | --- | --- | --- | --- | --- | --- | --- | --- |
| Region | I1 | I2a | I2b | J1 | J2a | J2b | R1a | R1b | G1 | G2a | E1b1b | Q | T | L |
| Hvar | 3 | 55 |  |  | 4 | 1 | 11 | 4 |  | 6 | 12 | 8 |  |  |
| Varaždin | 8 | 18 | 3 | 1 | 4 | 3 | 38 | 9 |  |  | 16 |  |  |  |
| Split | 5 | 53 | 2 |  | 2 | 3 | 20 | 5 | 1 |  | 11 | 1 | 2 |  |
| Pula | 9 | 31 | 1 | 1 | 3 | 2 | 28 | 8 |  | 3 | 11 |  | 1 | 1 |
| Osijek | 12 | 45 | 1 | 1 | 5 | 2 | 29 | 7 |  | 1 | 6 | 1 |  |  |
| Total number of haplotypes | 37 | 202 | 7 | 3 | 18 | 11 | 126 | 33 | 1 | 10 | 56 | 10 | 3 | 1 |

**FIGURE 4.**
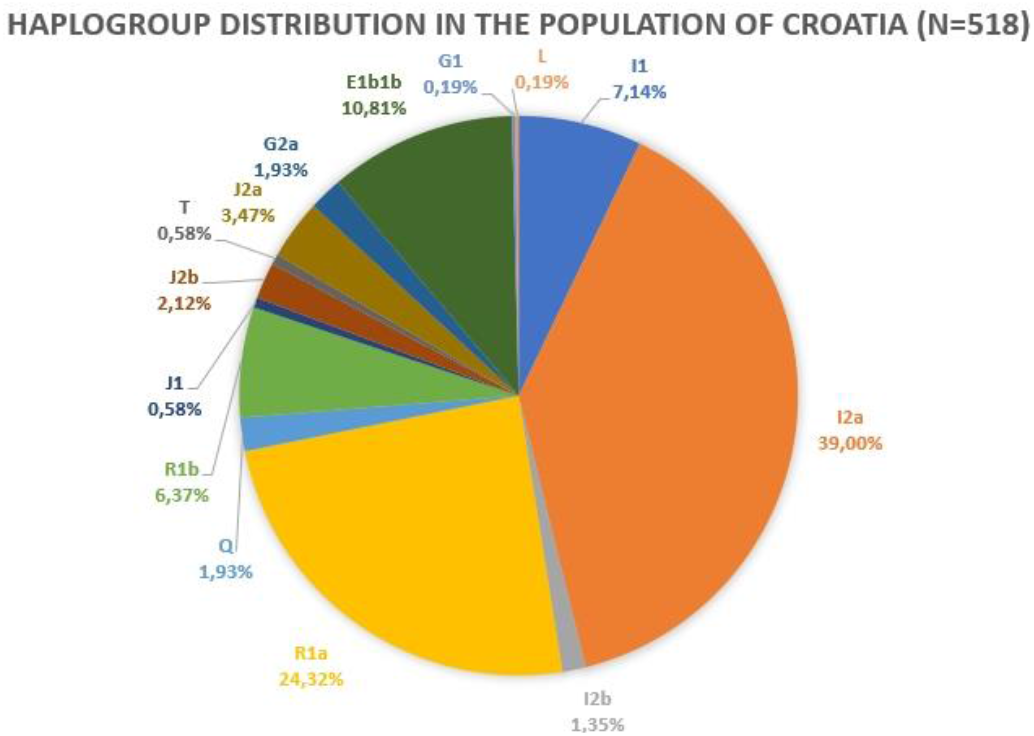
Y chromosome haplogroup prediction in the Croatian population (n=518) based on the population study of 27 Y-STR markers (Yfiler™ Plus PCR Amplification Kit, Applied Biosystems). The study included five Croatian regional subpopulations with the local centers in Osijek, Pula, Varaždin, Split, and Hvar Island

Four of the five subpopulations of Croatia showed expected results (Figure 5a-c). High frequency of haplogroup I have been reported with its known sublineage I2a in the subpopulations Hvar 52,88%, Split 50,48%, Osijek 40,91%, and Pula 31,31%. Previously published reports demonstrate similar results (12, 13, 16, 18, 19). However, slightly different results were obtained in the subpopulation of Varaždin (Figure 5d). R1a was the most frequent haplogroup in this subpopulation with a frequency of 38%, while the frequency for I2a haplogroup was 18%. Interestingly, R1a was also the dominant haplogroup within the Slovenian population (31), which is the closest neighboring abroad population to Varaždin county. However, as we have already stated, these initially notified similarities still should be confirmed by additional analysis and increasing/structuring sample size of the Varaždin region.

**FIGURE 5.**
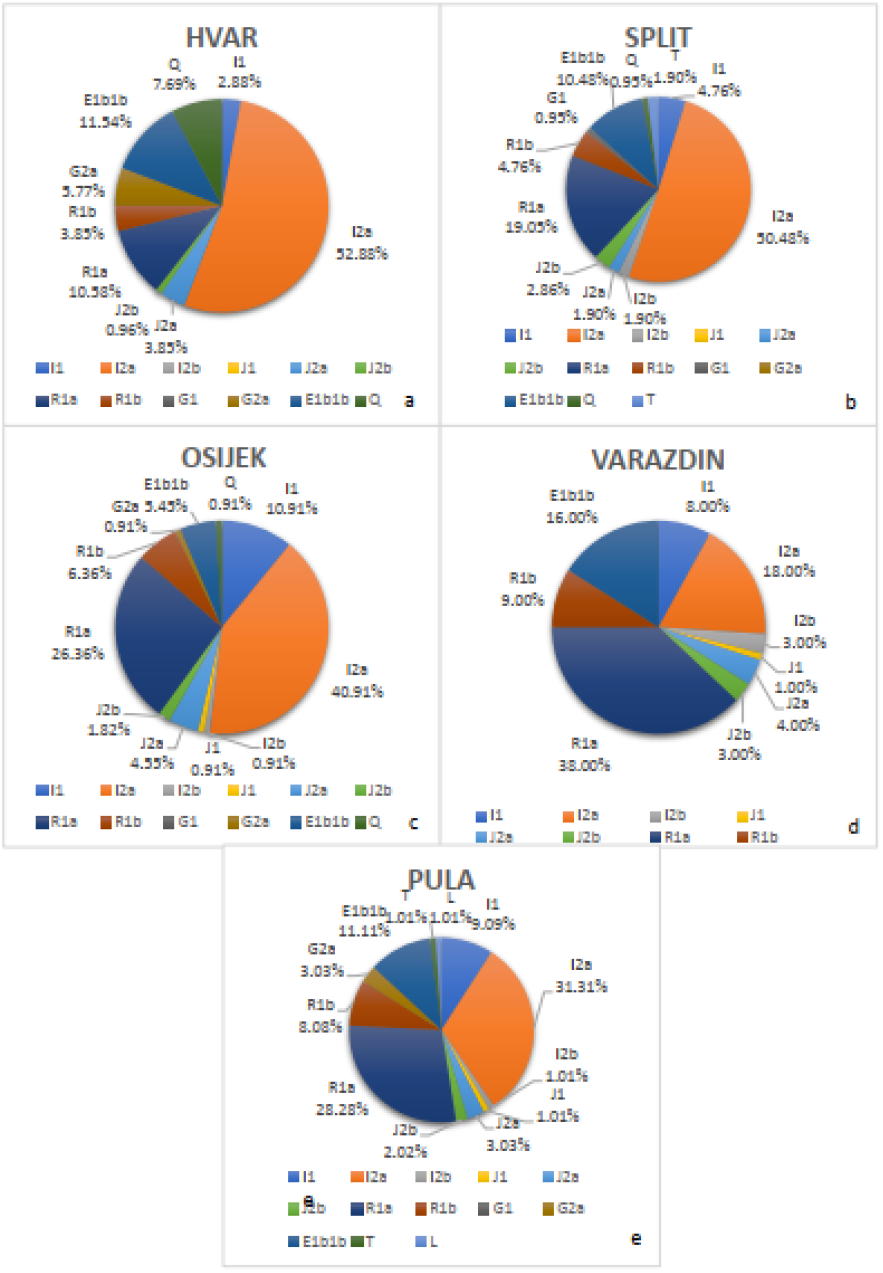
Y chromosome haplogroup frequency in five Croatian subpopulations: a. Hvar (n=104), b. Split (n=105), c. Osijek (n=110), d. Varaždin (n=100), e. Pula (n=99). Y haplogroup frequencies are determined in the population study of 27 Y-STR markers (Yfiler™ Plus PCR Amplification Kit, Applied Biosystems).

In summary, sublineage I2a was generally the most frequent haplogroup in the populations of Croatia, according to this study, but also to all previously studied data (12, 13, 16, 18, 19, 34). It is important to emphasize that similar results were obtained in an earlier study when I-P37 (a former name for the most I2a sublineage) for the Croatian population in Bosnia and Herzegovina was detected in a ratio of 71,1% (15).

Arrival of haplogroup I (previously described as Eu7 (12)) in this part of Europe is approximated to around 25,000 years ago from the Middle East through Anatolia in the area of the Balkan (9, 16). One scenario suggests the possibility of population expansion from one of the post-Glacial refuges into the rest of the Balkan Peninsula (15). There is also another possibility that this haplogroup could be connected with more recent population movements from Eastern Europe, but this idea still has to be examined (9). Definitively, when compared to the other populations in Europe, the I2a haplogroup sublineage is considered the characteristic Southeast European haplogroup (33).

The R1a (previously described as Eu 19 (12)) as a leading sublineage of haplogroup R was the second most frequent haplogroup in the studied population of Croatia, with an overall frequency of 24,32%. The prevalence of haplogroup R1a in the subpopulations of Croatia reported frequencies for Varaždin 38%, Pula 28,28%, Osijek 26,36% and Split 19,05%. In the subpopulation of Hvar, a small genetic deviation in the frequency of haplogroups R1a and E1b1b was reported. The R1a haplogroup accounted for 10,58% just slightly lower than haplogroup E1b1b with the frequency of 11,54% (Figure 5a). This is most likely due to the founder effect which is expected for island populations. In previously reported studies on the mainland population of Croatia, haplogroup R was reported as the second most frequent (13, 18, 34). Migration theories of R1a origins indicate the outflow of haplogroup R from West Asia to the region of Balkan as a post-Last Glacial Maximum (LGM) event during the Mesolithic time (16, 35).

Sublineage R1b (previously described as Eu 18 (12)) presented a lower frequency in the studied population of Croatia. The overall frequency of the R1b sublineage for the population of Croatia accounts of 6,37%. The highest frequency of R1b haplogroup was reported in the subpopulation of Varaždin with a prevalence of 9% and in Pula with a frequency of 8,08%. The most similar results were obtained in the Bosnian population based on 481 Y-STR profiles, whereby R1b accounted for 8,75% of the samples (10).

Sublineage E1b1b (previously predominantly described as Eu 4 (12)) is the most frequent “neolithic haplogroup” for the males in this part of Europe (16). In the present study E1b1b was detected with a frequency of 10,81%. The highest prevalence of this haplogroup was reported in the subpopulation of Varaždin, with a frequency of 16%. In the other four subpopulations the frequency of haplogroup E1b1b was estimated: Hvar 11,54%, Pula 11,11%, Split 10,48% and Osijek 5,45%. According to the recently published results, this haplogroup is slightly less frequent than in the closest neighboring population of Bosnia and Herzegovina (14,58%) (10). There are two suggested theories about E1b1b arrival in Europe. One theory is a post-LGM event from Asia and Africa during Neolithic period of time, while the other evidence suggests that this haplogroup is Balkan-specific, and originated around 8,000 years ago as a part of Greek colonization among the northern part of the Peninsula (16, 32). This ancient European haplogroup shows its possible dual origin from two different source populations, during the recolonization of Europe from Iberia and from West Asia (16, 34).

An approximate comparison between frequencies of the earlier used Y chromosome lineage (Eu) determined by Semino et al. (12) and the frequency of the currently used haplogroups detected in the Croatian population is shown in Table 3. It is important to emphasize that exact comparison is not possible because current nomenclature offers more detailed and precise insight into Y chromosome diversity. However, this table could approximate a comparison between early and currently detected Y chromosome diversity within the Croatian population.

**TABLE 3.** An approximate comparison between frequencies of the earlier used Y chromosome lineage (Eu) determined by Semino et al. (12) and the frequency of the currently used haplogroups detected in the Croatian population. The exact comparison is not possible because current nomenclature offers more detailed and precise insight into Y chromosome diversity.

| EU | Current Hg | Closest Joint Mutation | Frequencies (%) |  |
| --- | --- | --- | --- | --- |
|  |  |  | Semino et al. (year 2000) | Current Croatian data (year 2022) |
| EU 4 | E1b1b | M35 | 6,9 | 10,8 |
| EU 7 | I | M170 | 44,8 | 47,49 |
| EU 18 | R1b | M173 | 10,3 | 6,37 |
| EU 19 | R1a | M17 | 29,3 | 24,32 |

Rare haplogroups discovered in this study were Q, T, L and G1 each present in 1,93%, 0,58%, 0,19% and 0,19% of all samples, respectively. Haplogroup L is associated with South Asia and India but is also found in low frequencies in Central Asia, Southwest Asia, and Southern Europe. With its alternative phylogenetic name K1a, haplogroup L is closely related to haplogroup T (36). Haplogroup T, also known by its phylogenetic name K1b, possibly originated in Western Asia and spread to East Africa, South Asia, and Southern Europe (37, 38). Finally, haplogroup Q represents the only Pan-American haplogroup and confirms the Asian origin of Native Americans, and provides insight into the main Asian-American migrations (39). Haplogroup G1 is detected for the very first time within the Croatian population. This haplogroup is found predominantly in the Eurasian population, particularly in Iran, and is very rare in Europe. Some authors suggest that this rare haplogroup could have been carried by the expansion of Iranian speakers northwards to the Eurasian steppe (40). However, its origin is still not clearly described.

The present analyses generally confirmed previous results and provided a more detailed insight into the genetic diversity of the subpopulations of Croatia. Analysis of 27 Y-STR loci on the currently studied population of Croatia was quantified through *R*st values and calculated based on the results of the same set of markers from other populations from the YHRD. The results indicate that the Croatian population does not deviate significantly from the neighboring populations of Bosnia and Herzegovina, Slovenia, and Serbia. This proves that the Y chromosome genetic marker has a noticeable geographical background (2, 15), and this analysis resulted in expected geographic clustering.

A previous study (16) concluded that most of the Croatian men (“owners” of HgI, R1a and R1b) are harboring the ancestral genetic impact of Old European people who settled in Europe approximately 25.000 – 30.000 years ago and survived the LGM in several different refugia. Results of our study on new additional Y-STR loci confirmed that more than 78% of the contemporary Croatians are included in that group. The rest of the population relates to the people who arrived mostly during the Neolithization process. A small portion of examined population originated from the “owners” of rare haplogroups in the term of European genetic diversity, of which the origin is still not clarified. Usage of additional Y-STR loci revealed detailed insight and supplementary information within the highly diverse and significant genetic diversity of the Croatian population.

## Acknowledgment

The authors thank the sample donors for participating in this research, and colleagues at the University Hospital Center, Split, Croatia; University Department of Forensic Sciences, University of Split, Split, Croatia; University North, Varaždin, Croatia; Faculty of Medicine, University of Osijek, Osijek, Croatia; Faculty of Dental Medicine and Health, University of Osijek, Osijek Croatia; School of Medicine Rijeka, University of Rijeka, Croatia; and General Hospital Pula, Pula, Croatia for assistance with the buccal swab collection.

This collaboration was supported by the Croatian Society for Human Genetics and the International Society for Applied Biological Sciences (ISABS).

## Ethical approval

was given by the Ethics Committee, Institute for Anthropological Research, Zagreb, Croatia (approval number 20211053).

---

**SUPPLEMENTARY TABLE 1.** Interpopulation comparison over 17 Y-STR loci (included in the Yfiler™ marker set, Applied Biosystems) of the current data with 20 previously published European populations using genetic distance *R*st values and *P* values. Data was obtained from the relevant publications and accessed through the YHRD.

| Population | Croatia | Austria | Belgium | Bosnia and Herzegovina | Bulgaria | Germany | Greece | Hungary | Italy | Poland | Serbia | Albania | Czech Republic | Estonia | Ireland | Lithuania | North Macedonia | Norway | Slovenia | Sweden | Ukraine |
| --- | --- | --- | --- | --- | --- | --- | --- | --- | --- | --- | --- | --- | --- | --- | --- | --- | --- | --- | --- | --- | --- |
| Croatia | - | 0.0000 | 0.0000 | 0.0002 | 0.0089 | 0.0000 | 0.0000 | 0.0000 | 0.0000 | 0.0000 | 0.0000 | 0.0000 | 0.0000 | 0.0000 | 0.0000 | 0.0000 | 0.0000 | 0.0000 | 0.0000 | 0.0000 | 0.0001 |
| Austria | 0.0795 | - | 0.0000 | 0.0000 | 0.0000 | 0.0000 | 0.0000 | 0.0000 | 0.0035 | 0.0000 | 0.0000 | 0.0000 | 0.0079 | 0.0000 | 0.0000 | 0.0000 | 0.0000 | 0.0000 | 0.0000 | 0.0000 | 0.0000 |
| Belgium | 0.1429 | 0.0188 | - | 0.0000 | 0.0000 | 0.0000 | 0.0000 | 0.0000 | 0.0000 | 0.0000 | 0.0000 | 0.0000 | 0.0007 | 0.0000 | 0.0000 | 0.0000 | 0.0000 | 0.0000 | 0.0000 | 0.0000 | 0.0000 |
| Bosnia and Herzegovina | 0.0076 | 0.0988 | 0.1671 | - | 0.0034 | 0.0000 | 0.0000 | 0.0000 | 0.0000 | 0.0000 | 0.0002 | 0.0000 | 0.0000 | 0.0000 | 0.0000 | 0.0000 | 0.0000 | 0.0000 | 0.0000 | 0.0000 | 0.0000 |
| Bulgaria | 0.0144 | 0.0649 | 0.1467 | 0.0186 | - | 0.0000 | 0.1307 | 0.0001 | 0.0000 | 0.0000 | 0.0095 | 0.0041 | 0.0000 | 0.0000 | 0.0000 | 0.0000 | 0.0706 | 0.0000 | 0.0001 | 0.0000 | 0.0003 |
| Germany | 0.1036 | 0.0214 | 0.0173 | 0.1382 | 0.1206 | - | 0.0000 | 0.0000 | 0.0000 | 0.0000 | 0.0000 | 0.0000 | 0.2638 | 0.0000 | 0.0000 | 0.0000 | 0.0000 | 0.0000 | 0.0000 | 0.0000 | 0.0000 |
| Greece | 0.0241 | 0.0519 | 0.1187 | 0.0269 | 0.0048 | 0.1025 | - | 0.0001 | 0.0008 | 0.0000 | 0.0001 | 0.0024 | 0.0000 | 0.0000 | 0.0000 | 0.0000 | 0.0785 | 0.0000 | 0.0000 | 0.0000 | 0.0000 |
| Hungary | 0.0238 | 0.0252 | 0.0623 | 0.0451 | 0.0354 | 0.0321 | 0.0283 | - | 0.0001 | 0.0000 | 0.0000 | 0.0000 | 0.0024 | 0.0000 | 0.0000 | 0.0000 | 0.0000 | 0.0000 | 0.0973 | 0.0000 | 0.0130 |
| Italy | 0.0659 | 0.0144 | 0.0441 | 0.0744 | 0.0465 | 0.0546 | 0.0223 | 0.0273 | - | 0.0000 | 0.0000 | 0.0000 | 0.0000 | 0.0000 | 0.0000 | 0.0000 | 0.0000 | 0.0000 | 0.0000 | 0.0000 | 0.0000 |
| Poland | 0.1216 | 0.1049 | 0.1305 | 0.1687 | 0.1603 | 0.0580 | 0.1555 | 0.0679 | 0.1542 | - | 0.0000 | 0.0000 | 0.0000 | 0.0000 | 0.0000 | 0.0000 | 0.0000 | 0.0000 | 0.0000 | 0.0000 | 0.0000 |
| Serbia | 0.0186 | 0.1087 | 0.1870 | 0.0068 | 0.0145 | 0.1612 | 0.0182 | 0.0612 | 0.0735 | 0.2021 | - | 0.0000 | 0.0000 | 0.0000 | 0.0000 | 0.0000 | 0.0000 | 0.0000 | 0.0000 | 0.0000 | 0.0000 |
| Albania | 0.0728 | 0.0960 | 0.1714 | 0.0585 | 0.0258 | 0.1723 | 0.0159 | 0.0865 | 0.0527 | 0.2372 | 0.0336 | - | 0.0000 | 0.0000 | 0.0000 | 0.0000 | 0.0112 | 0.0000 | 0.0000 | 0.0000 | 0.0000 |
| Czech Republic | 0.0767 | 0.0156 | 0.0252 | 0.1086 | 0.0875 | 0.0012 | 0.0704 | 0.0176 | 0.0390 | 0.0519 | 0.1330 | 0.1324 | - | 0.0000 | 0.0000 | 0.0000 | 0.0000 | 0.0018 | 0.0002 | 0.0000 | 0.0000 |
| Estonia | 0.1877 | 0.1038 | 0.0940 | 0.2198 | 0.2054 | 0.0623 | 0.1630 | 0.1043 | 0.1226 | 0.1052 | 0.2477 | 0.2263 | 0.0693 | - | 0.0000 | 0.0080 | 0.0000 | 0.0000 | 0.0000 | 0.0000 | 0.0000 |
| Ireland | 0.3178 | 0.1360 | 0.0852 | 0.3403 | 0.3446 | 0.1221 | 0.3169 | 0.2276 | 0.2137 | 0.2475 | 0.3677 | 0.3590 | 0.1671 | 0.2125 | - | 0.0000 | 0.0000 | 0.0000 | 0.0000 | 0.0000 | 0.0000 |
| Lithuania | 0.1706 | 0.1162 | 0.1109 | 0.2114 | 0.2008 | 0.0615 | 0.1754 | 0.0977 | 0.1484 | 0.0555 | 0.2415 | 0.2548 | 0.0652 | 0.0121 | 0.2139 | - | 0.0000 | 0.0000 | 0.0000 | 0.0000 | 0.0000 |
| North Macedonia | 0.0375 | 0.0747 | 0.1458 | 0.0312 | 0.0067 | 0.1358 | 0.0036 | 0.0528 | 0.0419 | 0.1887 | 0.0165 | 0.0067 | 0.1028 | 0.2029 | 0.3271 | 0.2178 | - | 0.0000 | 0.0000 | 0.0000 | 0.0000 |
| Norway | 0.1270 | 0.0391 | 0.0394 | 0.1616 | 0.1473 | 0.0160 | 0.1183 | 0.0500 | 0.0715 | 0.0862 | 0.1797 | 0.1913 | 0.0178 | 0.0553 | 0.1700 | 0.0670 | 0.1535 | - | 0.0000 | 0.0000 | 0.0000 |
| Slovenia | 0.0204 | 0.0376 | 0.0887 | 0.0463 | 0.0297 | 0.0466 | 0.0284 | 0.0025 | 0.0439 | 0.0639 | 0.0606 | 0.0868 | 0.0269 | 0.1196 | 0.2641 | 0.1046 | 0.0511 | 0.0589 | - | 0.0000 | 0.0281 |
| Sweden | 0.1209 | 0.0427 | 0.0447 | 0.1437 | 0.1301 | 0.0400 | 0.0894 | 0.0545 | 0.0449 | 0.1360 | 0.1546 | 0.1406 | 0.0363 | 0.0646 | 0.2141 | 0.0956 | 0.1176 | 0.0147 | 0.0674 | - | 0.0000 |
| Ukraine | 0.0195 | 0.0603 | 0.1135 | 0.0461 | 0.0405 | 0.0599 | 0.0384 | 0.0079 | 0.0619 | 0.0530 | 0.0640 | 0.1005 | 0.0396 | 0.1254 | 0.2892 | 0.1021 | 0.0632 | 0.0845 | 0.0060 | 0.0950 | - |

**SUPPLEMENTARY TABLE 2.** Interpopulation comparison over 27 Y-STR loci (included in the Yfiler™ Plus PCR Amplification Kit, Applied Biosystems) of the current data with 23 previously published Worldwide populations using genetic distance *R*st values and *P* values. Data was obtained from the relevant publications and accessed through the YHRD.

| Population | Croatia | Austria | Belgium | Denmark | Ethiopia | French Polynesia | Germany | Ghana | Hungary | India | Italy | Lithuania | Mexico | Nigeria | North Macedonia | Pakistan | Poland | Russian Federation | Saudi Arabia | Serbia | Slovenia | Spain | Switzerland | United Kingdom |
| --- | --- | --- | --- | --- | --- | --- | --- | --- | --- | --- | --- | --- | --- | --- | --- | --- | --- | --- | --- | --- | --- | --- | --- | --- |
| Croatia | - | 0.0000 | 0.0000 | 0.0000 | 0.0000 | 0.0000 | 0.0000 | 0.0000 | 0.0000 | 0.0000 | 0.0000 | 0.0000 | 0.0000 | 0.0000 | 0.0000 | 0.0000 | 0.0000 | 0.0000 | 0.0000 | 0.0055 | 0.0000 | 0.0000 | 0.0000 | 0.0000 |
| Austria | 0.1255 | - | 0.0000 | 0.0024 | 0.0000 | 0.0000 | 0.0052 | 0.0000 | 0.0000 | 0.0000 | 0.0000 | 0.0000 | 0.0000 | 0.0000 | 0.0000 | 0.0000 | 0.0000 | 0.0000 | 0.0000 | 0.0000 | 0.0000 | 0.0000 | 0.0000 | 0.0000 |
| Belgium | 0.2031 | 0.0197 | - | 0.0000 | 0.0000 | 0.0000 | 0.0027 | 0.0000 | 0.0000 | 0.0000 | 0.0000 | 0.0000 | 0.0000 | 0.0000 | 0.0000 | 0.0000 | 0.0000 | 0.0000 | 0.0000 | 0.0000 | 0.0000 | 0.0000 | 0.4444 | 0.0000 |
| Denmark | 0.1290 | 0.0099 | 0.0412 | - | 0.0000 | 0.0000 | 0.0013 | 0.0000 | 0.0000 | 0.0000 | 0.0000 | 0.0000 | 0.0000 | 0.0000 | 0.0000 | 0.0000 | 0.0000 | 0.0000 | 0.0000 | 0.0000 | 0.0000 | 0.0000 | 0.0000 | 0.0000 |
| Ethiopia | 0.1704 | 0.2165 | 0.2801 | 0.2495 | - | 0.0000 | 0.0000 | 0.0000 | 0.0000 | 0.0000 | 0.0000 | 0.0000 | 0.0000 | 0.0000 | 0.0000 | 0.0000 | 0.0000 | 0.0000 | 0.0000 | 0.0000 | 0.0000 | 0.0000 | 0.0000 | 0.0000 |
| French Polynesia | 0.1878 | 0.1419 | 0.2024 | 0.1934 | 0.2763 | - | 0.0000 | 0.0000 | 0.0000 | 0.0000 | 0.0000 | 0.0000 | 0.0000 | 0.0000 | 0.0000 | 0.0000 | 0.0000 | 0.0000 | 0.0000 | 0.0000 | 0.0000 | 0.0000 | 0.0000 | 0.0000 |
| Germany | 0.1461 | 0.0045 | 0.0104 | 0.0102 | 0.2501 | 0.1709 | - | 0.0000 | 0.0000 | 0.0000 | 0.0000 | 0.0000 | 0.0000 | 0.0000 | 0.0000 | 0.0000 | 0.0000 | 0.0000 | 0.0000 | 0.0000 | 0.0000 | 0.0000 | 0.0000 | 0.0000 |
| Ghana | 0.2787 | 0.3710 | 0.4691 | 0.4133 | 0.2459 | 0.3321 | 0.4094 | - | 0.0000 | 0.0000 | 0.0000 | 0.0000 | 0.0000 | 0.0012 | 0.0000 | 0.0000 | 0.0000 | 0.0000 | 0.0000 | 0.0000 | 0.0000 | 0.0000 | 0.0000 | 0.0000 |
| Hungary | 0.0482 | 0.0757 | 0.1484 | 0.0879 | 0.1303 | 0.1706 | 0.0947 | 0.3044 | - | 0.0000 | 0.0000 | 0.0000 | 0.0000 | 0.0000 | 0.0000 | 0.0000 | 0.0000 | 0.0000 | 0.0000 | 0.0000 | 0.0004 | 0.0000 | 0.0000 | 0.0000 |
| India | 0.0961 | 0.1023 | 0.1642 | 0.1154 | 0.1454 | 0.1681 | 0.1217 | 0.2779 | 0.0201 | - | 0.0000 | 0.0000 | 0.0000 | 0.0000 | 0.0000 | 0.0000 | 0.0000 | 0.0000 | 0.0000 | 0.0000 | 0.0000 | 0.0000 | 0.0000 | 0.0000 |
| Italy | 0.2043 | 0.0249 | 0.0159 | 0.0575 | 0.2656 | 0.1578 | 0.0281 | 0.4213 | 0.1339 | 0.1529 | - | 0.0000 | 0.0000 | 0.0000 | 0.0000 | 0.0000 | 0.0000 | 0.0000 | 0.0000 | 0.0000 | 0.0000 | 0.0000 | 0.0000 | 0.0000 |
| Lithuania | 0.1914 | 0.1028 | 0.1330 | 0.0847 | 0.2836 | 0.2703 | 0.0922 | 0.4656 | 0.1106 | 0.1226 | 0.1454 | - | 0.0000 | 0.0000 | 0.0000 | 0.0000 | 0.0000 | 0.0000 | 0.0000 | 0.0000 | 0.0000 | 0.0000 | 0.0000 | 0.0000 |
| Mexico | 0.1785 | 0.0949 | 0.1173 | 0.1287 | 0.2192 | 0.1469 | 0.1094 | 0.3530 | 0.1398 | 0.1410 | 0.1291 | 0.2279 | - | 0.0000 | 0.0000 | 0.0000 | 0.0000 | 0.0000 | 0.0000 | 0.0000 | 0.0000 | 0.0000 | 0.0000 | 0.0000 |
| Nigeria | 0.2414 | 0.3309 | 0.4230 | 0.3709 | 0.2023 | 0.2829 | 0.3721 | 0.0054 | 0.2563 | 0.2394 | 0.3853 | 0.4213 | 0.3107 | - | 0.0000 | 0.0000 | 0.0000 | 0.0000 | 0.0000 | 0.0000 | 0.0000 | 0.0000 | 0.0000 | 0.0000 |
| North Macedonia | 0.0720 | 0.0374 | 0.0848 | 0.0627 | 0.1212 | 0.1627 | 0.0548 | 0.3156 | 0.0348 | 0.0656 | 0.0780 | 0.1377 | 0.1012 | 0.2687 | - | 0.0000 | 0.0000 | 0.0000 | 0.0000 | 0.0000 | 0.0000 | 0.0000 | 0.0000 | 0.0000 |
| Pakistan | 0.0854 | 0.0531 | 0.1030 | 0.0519 | 0.2216 | 0.1213 | 0.0644 | 0.3449 | 0.0381 | 0.0503 | 0.1003 | 0.0963 | 0.1256 | 0.2998 | 0.0718 | - | 0.0000 | 0.0000 | 0.0000 | 0.0000 | 0.0000 | 0.0000 | 0.0000 | 0.0000 |
| Poland | 0.0905 | 0.1294 | 0.1886 | 0.1076 | 0.2569 | 0.2681 | 0.1225 | 0.4045 | 0.0687 | 0.0972 | 0.2007 | 0.0684 | 0.2363 | 0.3703 | 0.1198 | 0.0762 | - | 0.0000 | 0.0000 | 0.0000 | 0.0000 | 0.0000 | 0.0000 | 0.0000 |
| Russian Federation | 0.0779 | 0.0677 | 0.1188 | 0.0648 | 0.1586 | 0.1833 | 0.0794 | 0.2947 | 0.0303 | 0.0593 | 0.1229 | 0.0704 | 0.1350 | 0.2609 | 0.0471 | 0.0689 | 0.0635 | - | 0.0000 | 0.0000 | 0.0000 | 0.0000 | 0.0000 | 0.0000 |
| Saudi Arabia | 0.0990 | 0.1150 | 0.1891 | 0.1464 | 0.0732 | 0.1506 | 0.1523 | 0.2084 | 0.0562 | 0.0604 | 0.1695 | 0.2070 | 0.0956 | 0.1639 | 0.0577 | 0.1029 | 0.1896 | 0.0794 | - | 0.0000 | 0.0000 | 0.0000 | 0.0000 | 0.0002 |
| Serbia | 0.0097 | 0.1259 | 0.2101 | 0.1415 | 0.1231 | 0.2006 | 0.1525 | 0.2604 | 0.0449 | 0.0847 | 0.2057 | 0.2023 | 0.1647 | 0.2159 | 0.0517 | 0.1014 | 0.1189 | 0.0743 | 0.0669 | - | 0.0000 | 0.0000 | 0.0000 | 0.0000 |
| Slovenia | 0.0297 | 0.0671 | 0.1355 | 0.0615 | 0.1703 | 0.1821 | 0.0781 | 0.3378 | 0.0142 | 0.0514 | 0.1387 | 0.0932 | 0.1533 | 0.2887 | 0.0418 | 0.0334 | 0.0310 | 0.0275 | 0.0899 | 0.0423 | - | 0.0000 | 0.0000 | 0.0000 |
| Spain | 0.3283 | 0.1183 | 0.0490 | 0.1695 | 0.4077 | 0.3475 | 0.0887 | 0.5812 | 0.2931 | 0.2865 | 0.0760 | 0.2378 | 0.2156 | 0.5509 | 0.2032 | 0.2321 | 0.2990 | 0.2268 | 0.3483 | 0.3590 | 0.2795 | - | 0.0000 | 0.0000 |
| Switzerland | 0.1999 | 0.0138 | -0.0003 | 0.0299 | 0.2845 | 0.1833 | 0.0075 | 0.4310 | 0.1421 | 0.1632 | 0.0180 | 0.1283 | 0.1143 | 0.3997 | 0.0840 | 0.1026 | 0.1816 | 0.1170 | 0.1850 | 0.2052 | 0.1314 | 0.0581 | - | 0.0000 |
| United Kingdom | 0.1122 | 0.0813 | 0.1442 | 0.1122 | 0.0780 | 0.1429 | 0.1167 | 0.2236 | 0.0562 | 0.0558 | 0.1344 | 0.1718 | 0.0994 | 0.1752 | 0.0438 | 0.0865 | 0.1744 | 0.0756 | 0.0181 | 0.0808 | 0.0809 | 0.3041 | 0.1438 | - |

